# A cell-based method to assess the activity of the human sodium-coupled citrate transporter

**DOI:** 10.1101/2025.08.02.668312

**Authors:** Natali Cárcamo-Lemus, Angelo Bernier, Ivana Oporto-Ortega, Alejandro San Martín, Pamela Y. Sandoval

## Abstract

Increasing evidence supports the human sodium-coupled citrate transporter (hNaCT) as a potential therapeutic target for metabolic syndrome, while genetic polymorphisms in hNaCT have been associated with neurological disorder. Isotopic tracers remain the primary method for evaluating citrate transport, which are labor-intensive, require dedicated equipment, and demand highly skilled personnel. Herein, we develop and validate a robust, live-cell-compatible fluorescent method to evaluate citrate transport activity mediated by hNaCT at single-cell and high-throughput levels. This method utilizes a baculoviral vector to modify HEK293 cells to co-express a genetically encoded citrate sensor (Citron1) and the hNaCT. This cell-based platform enabled real-time monitoring of citrate transport using fluorescent microscopy and standard multiwell plate reader. The present method enables the functional characterization of hNaCT gain- and loss-of-function mutation, as well as the evaluation of pharmacological inhibitors targeting citrate transport. These results demonstrate that the cell-based method reliably assesses citrate transport at a single-cell resolution and is compatible with high-throughput screening, to identify novel lead compounds for the therapeutic modulation of hNaCT.

## 1 INTRODUCTION

Under physiological conditions, citrate is synthesized in the mitochondrial matrix as part of the tricarboxylic acid (TCA) cycle. During anabolic states, citrate is exported to the cytosol, where it supports fatty acid and cholesterol biosynthesis. Additionally, cytosolic citrate levels act as indicators of cellular energy status. For instance, low citrate concentrations stimulate glycolysis, the TCA cycle, and fatty acid oxidation (1, 2).

Cell surface citrate uptake is primarily mediated by the sodium-coupled citrate transporter (NaCT) in the brain and liver of mammals (3). This transporter has been mainly characterized by using isotopic methods. While these techniques are valuable, they lack single-cell resolution, as the averaging of signals across thousands of cells canceling-out cell-to-cell variability. Additionally, they destroy the sample, precluding before-after experimental design. New methods with single-cell resolution and easy-to-detect readouts would be valuable to unveil metabolic heterogeneity and could be readily adapted for high-throughput applications in pathological contexts.

Studies in flies have shown that deletion of NaCT mimics caloric restriction, which has been associated with increased life span (4). Furthermore, reducing NaCT expression leads to increased mitochondrial biogenesis, enhanced lipid oxidation, and improved insulin sensitivity in mice (5,6). Additionally, its polymorphism has been linked to congenital epilepsy (7). These findings suggest hNaCT as a therapeutic target for metabolic diseases and neurological disorders. Despite promising evidence suggesting potential benefits in developing modulator drugs, no hNaCT inhibitor has advanced to clinical trials (8), highlighting the need to expand the diversity of drugs targeting hNaCT. Radiolabeled [^14^C]-citrate uptake assays are the most used high-throughput approach for screening NaCT transport inhibitors. However, this technique remains limited in its scalability and adaptability for high-throughput screening formats.

This study introduces new methods to assess citrate transport at single-cell resolution and to establish a cell-based platform for high-throughput screening using the genetically encoded fluorescent sensor Citron1 (9). These approaches enable functional characterization of hNaCT at the single-cell level and support the implementation of a high-throughput, cell-based system for identifying citrate transporter modulators.

## 2 MATERIALS AND METHODS

### 2.1 Reagents

Trisodium citrate dihydrate (Sigma-Aldrich, CAS No.6132-04-3), lithium chloride (Sigma-Aldrich, CAS No.7447-41-8), and PF-06761281-compound-4a (Sigma-Aldrich, CAS No.1854061-19-0) were diluted in water. Phloretin (Sigma-Aldrich, CAS No.60-82-2) was prepared in 70% ethanol. BI01383298 (Sigma-Aldrich, CAS No.2227549-00-8) and 4-(Chloromercuri) benzenesulfonic acid sodium salt (pCMBS) (Toronto Research Chemicals, CAS No.14110-97-5) were prepared in 0.5 % DMSO. Unless limited by the maximum solubility indicated by the manufacturer, all inhibitors were prepared at 1000 times the working concentration.

### 2.2 Single-cell functional assay

HEK293 cells were obtained from the ATCC and cultured at 37 °C in 5% CO2 in Dulbecco’s modified Eagle’s medium/F-12 supplemented with 10% fetal bovine serum. For single-cell fluorescence microscopy assays, cells were seeded onto 25 mm sterile coverslips pretreated with 0.01% poly-L-lysine (Gibco, CAS No.25988-63-0). Transduction was done directly on cultures growing on the coverslip using a BacMam baculoviral vector system encoding Citron1 (Addgene #134303) and hNaCT (synthesized by Genscript). Alternatively, transfection of cells was done at 60% confluence using Lipofectamine3000 (ThermoFisher #L3000015) and studied after 16-24 h. Cells were superfused with KRH buffer of the following composition (in mM): 136 NaCl, 1.25 CaCl_2_, 1.25 MgSO_4_, 10 HEPES, 3 KCl, pH 7.4, using an upright Olympus FV1000-confocal microscope equipped with a 20× water immersion objective (N.A.1.0) and a 488 nm solid-state laser. Time series images were taken every 10s with 20× (NA.1.0) in XYT scan mode (scan speed: 156 Hz; 800 × 800-pixel; pinhole 800 μm). The fluorescent signal for a region of interest (ROI) from each cell was collected. Background subtraction was performed with ImageJ, and the statistical analysis was carried out using SigmaPlot (Jandel).

### 2.3 Multiwell plate assay

For culturing 96-well plates, wells were pre-treated with 0.01% poly-L-lysine. HEK293 cell suspension (250,000 cells/mL) with viability greater than 90% was inoculated with 30 μl/ml of baculoviral system (95% TU) for the co-expression of Citron1 (Addgene #134303) and hNaCT (synthesized by Genscript). Each well was seeded with 50,000 cells and incubated under appropriate conditions until maximal confluence was reached. Fluorescence was acquired using the Infinite F Nano+ microplate reader (Tecan). Cell tracker Calcein Red-Orange, AM (Invitrogen, Cat. No.C34851) was used to normalize the Citron1 fluorescence. Briefly, after the media removal, each well was washed twice with 200 μl of KRH buffer and incubated for 30 minutes. Then, the buffer was replaced with fresh KRH containing 1 μM Calcein Red-Orange and incubated at room temperature for 20 minutes. After removal of residual Calcein, 200 μl of KRH buffer was added, and the minimal fluorescence was measured in each well by sequential excitation at 485 nm (cpGFP) and 560 nm (Calcein Red-Orange) to record fluorescence emissions at 510 nm and 610 nm, respectively. Afterward, the buffer was replaced with 200 μl of KRH buffer containing the inhibitors at their respective concentration. Additional wells without inhibitors were left to evaluate maximal citrate uptake. The plate was incubated for 30 minutes, and the buffer was replaced with KRH containing the same previous treatment supplemented with 0.1 mM citrate. Finally, the plate was incubated, and fluorescence was measured in each well for up to 60 minutes.

### 2.4 Data Processing and Statistical Analysis

Statistical analyses were performed using SigmaPlot-12 software (Jandel). For the before-after comparisons involving normally distributed data, paired t-tests were conducted. To compare citrate uptake among different hNaCT genetic variants, a non-parametric Kruskal-Wallis test was applied, and post hoc comparisons were performed using Dunn’s method to identify statistically significant differences between groups.

To evaluate the performance of the high-throughput screening assay, the Z’-factor coefficient was employed. (10) The Z’-factor is dimensionless and is calculated using the following equation.

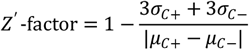

Where ơ is the standard deviation and µ is the mean of positive (C+) and negative (C-) controls. Z’-factor values between 0.5 and 1 indicate the ability to identify an inhibitor with 99.73% confidence using a single well. Values below 0.5 imply the need for duplicates or triplicates for each proposed inhibitor, and a Z’-factor below 0 indicates that the assay is unsuitable for high-throughput screening. The normalized percentage of inhibition (NPI) was calculated using the following equation to compare the transport inhibition induced by the tested drugs.

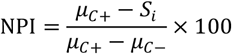

## 3 RESULTS

### 3.1 Single-cell citrate transport assay

To characterize hNaCT-mediated citrate transport at single-cell resolution, we established a heterologous expression system by co-expressing hNaCT and the highly responsive fluorescent citrate reporter Citron1 in HEK293 cells. These cell lines do not show endogenous citrate permeability, making it a perfect system to evaluate heterologous expressed citrate transporters. A baculoviral system was employed to drive the bicistronic expression of Citron1 with a version of hNaCT under the robust cytomegalovirus promoter. The correct localization pattern of mRuby3-tagged version of hNaCT in HEK293 cells was confirmed by fluorescence microscopy (**Figure 1A, top panel)**. The Citron1 molecular sensor exhibited a homogeneous expression pattern with a high signal-to-noise ratio, as originally reported (**Figure 1A, bottom panel**). Accordingly, our reporter system comprises HEK293 cells transduced with a baculoviral vector enabling the co-expression of Citron1 and hNaCT.

**Figure 1.**
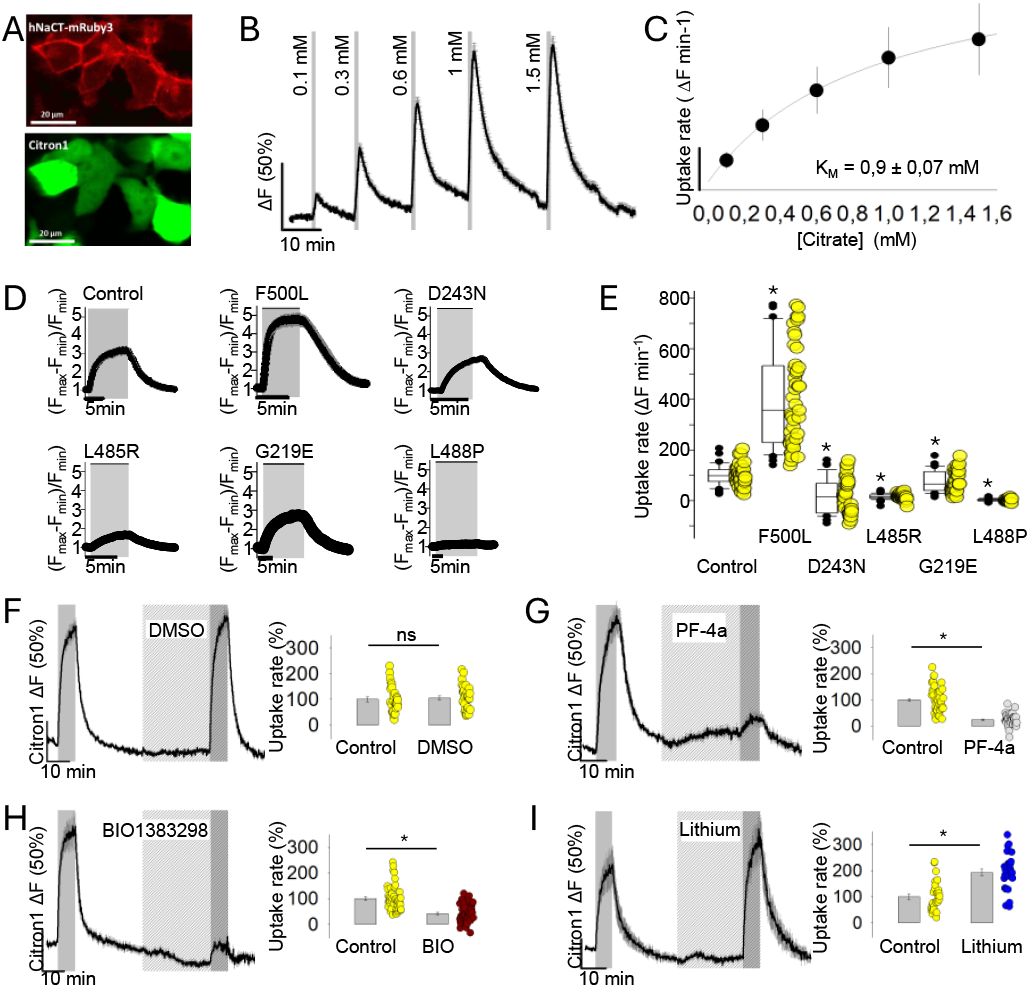
Single-cell fluorescence-based assay for monitoring hNACT-mediated citrate transport using Citron1. (**A**) Representative confocal images of HEK293 cell expressing hNaCT-mRuby3-tagged (top) and Citron1 (bottom), display predominant plasma membrane localization of hNaCT and cytosolic distribution of citrate sensor. Scale bar 20 µm. (**B**) Average fluorescence response of Citron1 from reporter cells exposed to increasing concentrations of extracellular citrate (0.1 mM to 1.5 mM). Fluorescence signals as mean ± SEM of 10 cells from a representative experiment. A total of 30 cells were analyzed across three independent experiments. (**C**) The affinity constant (K_m_) was determined by fitting the initial rate of citrate uptake, obtained from the cells described in panel B, to a rectangular hyperbola. The resulting mean ± SEM values were calculated from three independent experiments (n = 3, 30 cells total). (**D**) Recordings of Citron1 fluorescence signals following 0.1 mM citrate addition in cells expressing wild-type or the indicated variant versions of hNaCT. Each panel displays the mean ± SEM of 10 cells from representative experiments. (**E**) Functional comparison of hNaCT variants based on initial uptake rate. Box-and-whisker plots summarize single-cell uptake measurements (ΔF min^-1^) from reporter cells expressing either control hNaCT or the indicated variants (F500L, D243N, L485R, G219E, L488P). Each dot represents an individual cell; data was pooled from three independent experiments (30 cells total per variant). A non-parametric Kruskal-Wallis test was applied. (*) represent a p < 0.05 versus control. (**F-I**) Pharmacological modulation of hNaCT activity. Reporter cells were sequentially exposed to 0.1 mM citrate in absence and presence of: (**F**) vehicle control (0.5% DMSO), (**G**) 10 µM PF-4a, (**H**) 10 µM BIO1383298, and (**I**) 10mM lithium. Left panels: representative Citron1 fluorescence signals (mean ± SEM of 10 cells). Right panels: quantification of initial uptake rates from three independent experiments (n=3, 30 cells per condition). Data are shown as mean ± SEM. Paired t-test; ns = not significant; * = p < 0.05

Using our reporter cells, we designed a quantitative citrate transport method based on the uncalibrated fluorescence signal to characterize the citrate transporter. This quantitative method follows a before-and-after experimental design with comparable baseline intracellular levels prior to citrate addition. This methodology takes advantage of the first seconds of citrate uptake which are proportional to hNaCT activity and independent of Citron1 affinity. Using this approach the apparent K_m_ of hNaCT was determined. Reporter cells were exposed to increasing citrate concentrations (0.1 to 1.5 mM) under an open perfusion system in bicarbonate-free conditions, while intracellular citrate accumulation was monitored as a concentration-dependent increase in Citron1 fluorescence (**Figure 1B**). The initial rate of citrate uptake at each concentration was fitted to a rectangular hyperbola, yielding a K_m_ of 0.9 +/-0.07 mM (**Figure 1C**), which falls within the range of affinities previously reported for hNaCT (7,11). Using the same approach, we evaluated the sensitivity of the method to detect functional alteration in the hNaCT by analyzing well-characterized gain- and loss-of-function genetic variants, previously identified as naturally occurring mutations associated with congenital epilepsy (11). Loss-of-function mutations may result in reduced cell surface expression or impaired trafficking, while some variants retain proper localization but exhibit diminished activity. The response of the genetic variants to a rapid pulse of extracellular citrate showed uptake kinetics according to the published data associated with the polymorphisms (**Figure 1D**). We calculate the initial citrate uptake rate of all the genetic variants to validate our results (**Figure 1E**). Compared to the control, the gain-of-function F500L exhibited a significantly higher uptake rate (over 200%), consistent with enhanced transporter activity. In contrast, loss-of-function variants, such as D243N, L485R, and G219E, displayed a significantly diminished activity, while the L488P mutation resulted in a complete loss of citrate transport (**Figure 1E**). These results confirm that the citrate transport method allows us to evaluate the performance of hNaCT polymorphism associated with pathological conditions.

Next, we tested whether our method could detect pharmacological modulation of hNaCT activity using commercially available inhibitors and activators at single-cell resolution. Reporter cells were exposed to consecutive pulses of citrate 0.1 mM in absence and presence of a given treatment. To discard a potential solvent interference, we tested the citrate uptake in the presence of DMSO. The first citrate pulse was applied in buffer alone, while the second included DMSO. Comparison of the initial citrate uptake rates showed no significant differences between pulses, indicating that DMSO does not affect citrate transport (**Figure 1F**). Then, we assessed the inhibitory effects of PF-06761281 (hereafter referred to as PF-4a, (12)) and BI01383298 (13), two compounds previously characterized as potent and selective hNaCT inhibitors. Reporter cells were consecutively exposed to 0.1 mM citrate in the absence and subsequently in the presence of each inhibitor. As expected, both compounds induced a pronounced reduction in citrate uptake, PF-4a induced an average decrease of 80 % and BI01383298 achieving a 60 % reduction (**Figures 1G, H**). These findings indicate that the assay is both sensitive and robust enough to detect functional inhibition of hNaCT activity under the tested conditions. To further validate the responsiveness of our reporter system, we evaluated the well-documented enhancing effect of lithium on hNaCT-mediated citrate transport (14). Reporter cells incubated with 10 mM lithium exhibited a 1.5-fold increase in the initial rate of citrate uptake compared to untreated control conditions (**Figure 1I**). These results demonstrate that, under the current assay conditions, cells co-expressing hNaCT and Citron1 provide a reliable platform for the functional assessment of citrate transport, enabling the detection of both inhibitory and stimulatory modulation of hNaCT activity.

### 3.2 A fluorescence cell-based high-throughput method to assess citrate transport

Given the high dynamic range of Citron1, we developed a high-throughput cell-based assay for the functional evaluation of hNaCT in 96-well format. As proof of the concept, we used well-known hNaCT inhibitors such as PF-4a and BI01383298. Additionally, pCMBS was included as a non-specific yet potent inhibitor of transporter proteins due to its ability to modify sulfhydryl groups on cysteine residues. Phloretin, a broad-spectrum inhibitor known to interfere with various solute carriers, including hNaCT, was also evaluated as a non-selective reference compound. Our results demonstrated that the fluorescence signal remained stable across all treatment conditions for at least 60 minutes, confirming the temporal robustness of the assay (**Figure 2A**).

**Figure 2.**
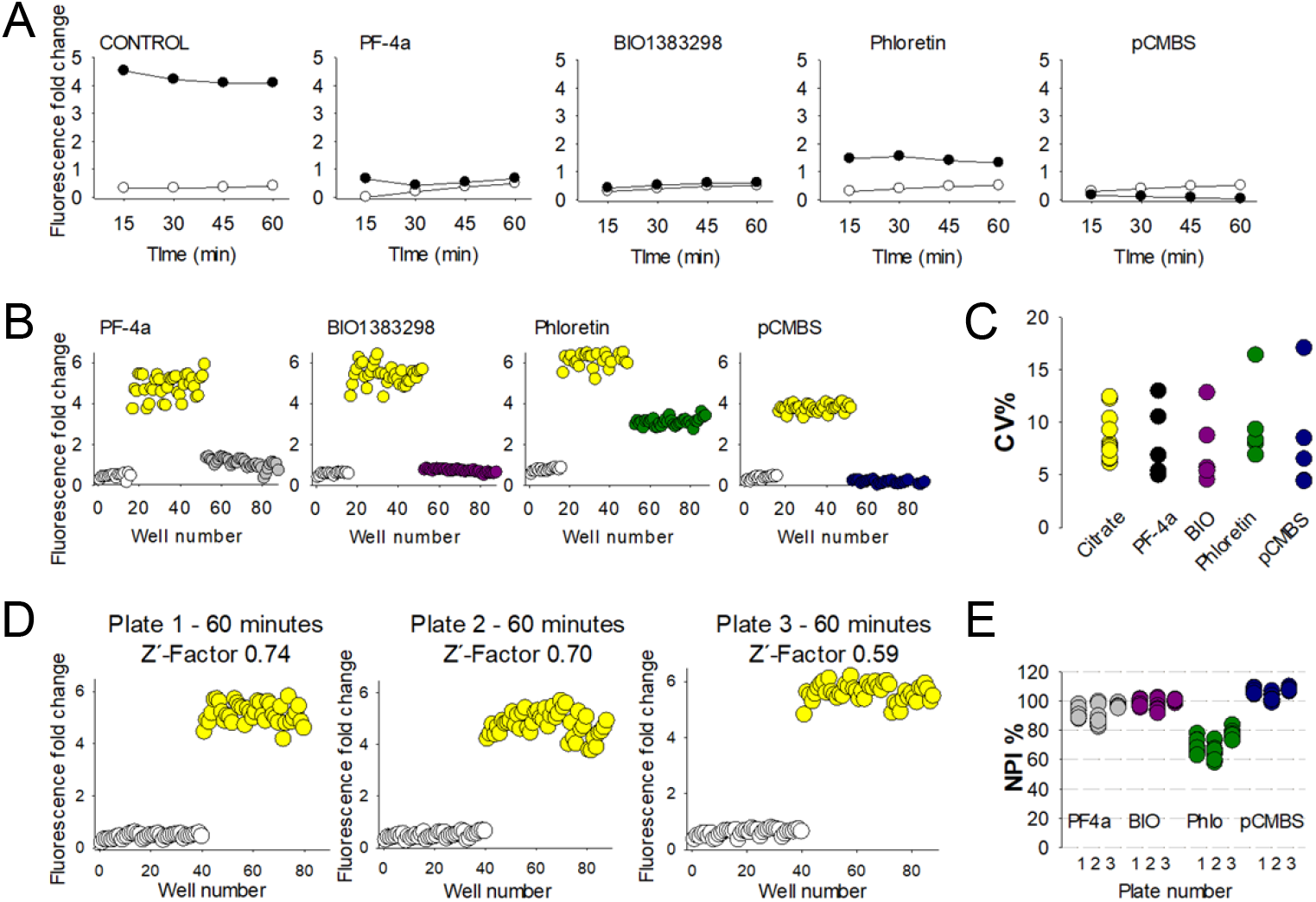
Validation of the cell-based high-throughput assay for hNaCT inhibition. (**A**) Time-course of fluorescence signal from reporter cells incubated with 0.1 mM citrate (filled circles) or vehicle (open circle), in the absence (control) or presence of known hNaCT inhibitors: 10 µM PF-06761281 (PF-4a), 10 µM BIO1383298, 100 µM phloretin or 10 µM PCMB. The fluorescence signal was monitored for over 60 minutes. Each panel represents the mean of at least 32 wells ± SEM. (**B**) Fluorescence output mapping across 96-well plates for each condition. Each dot represents an individual well. Yellow dots correspond to citrate-treated wells (positive control), reflecting maximal hNaCT-mediated transport activity. White dots represent vehicle-treated wells (negative control), while colored dots correspond to wells exposed to indicated hNaCT inhibitor. (**C**) The percentage of coefficient variation (CV%) across replicates for each treatment shows the inter-plate citrate uptake variation. (**D**) Z’-factor determination from three independent experiments at 60 minutes post treatment. Negative controls in white and positive controls in yellow (0.1 mM citrate). (**E**) Normalized percentage inhibition (NPI%) of citrate uptake across three biological replicates for each inhibitor. Each dot represents a single plate, bars denote mean ± SEM.

Consistent with their known inhibitory properties, PF-4a, BIO1383298, and pCMBS induced a marked reduction in the fluorescence signal associated with citrate uptake (**Figure 2B**). In contrast, phloretin displayed only a modest decrease in the fluorescent fold-change, reflecting its lower inhibitory efficacy (**Figure 2B**). Intra-plate variability, expressed as the coefficient of variation (CV%), was as low as 5 %, whereas inter-plate variability reached up to 18 %, indicating substantial variation between-plates (**Figure 2C**).

To evaluate the suitability of the assay for high-throughput applications, we calculated the Z’-factor, a statistical parameter that quantifies assay robustness and its ability to distinguish true positive hits from background variability (10). A Z’-factor values between 0.5 and 1 indicates excellent assay performance, allowing confident hit identification from single well measurements with 99.73 % - 100 % reliability. In three independent experiments comparing negative and positive controls at 60 minutes of treatment, the assay for citrate uptake mediated by hNaCT yielded Z’-factor values of 0.74, 0.7, and 0.59, demonstrating consistent and reliable performance across replicates (**Figure 2D**). These values indicate a reliable assay, with excellent dynamic separation between positive and negative controls, a critical requirement for effective HTS at single-well resolution. Finally, to quantify the extent of transport inhibition by each compound, the normalized percentage of inhibition (NPI%) was calculated across three independent experiments (**Figure 2E**). As anticipated, PF-4a, BI01383298, and pCMBS exhibited markedly greater inhibitory effects on hNaCT-mediated citrate uptake compared to phloretin, which showed only partial inhibition under the assay conditions.

## 4. DISCUSSION

In this study, we developed and validated a fluorescence-based assay for the evaluation of citrate transport activity mediated by the human sodium-coupled citrate transporter (hNaCT). By combining heterologous expression of hNaCT with the genetically encoded citrate sensor Citron1, we established a robust and scalable cell-based platform capable of reporting genetic or pharmacological perturbation of intracellular citrate accumulation in live HEK293 cells. First, we showed a method to obtain quantitative kinetic information using uncalibrated Citron1 readout at single-cell level. Additionally, 96-well plate assay was established and validated with known hNaCT citrate transporter inhibitors.

We applied the assay to hNaCT genetic variants to establish proof-of-concept assay. According to previous report (11) the gain-of-function mutant F500L displayed an initial rate of citrate uptake over 200% higher than that observed in the hNaCT control condition. In contrast, loss-of-function mutants, such as D243N, L485R, and G219E, displayed a significantly diminished, or completely abolished in L488P, citrate uptake. These confirm that the platform successfully distinguished between gain- and loss-of-function mutations based on their impact on citrate transport, without apparent interference in Citron1 expression or functionality. This illustrates the flexibility of the system for assessing the functional consequences of hNaCT polymorphisms. Our results using Citron1 to characterize alterations on hNaCT activity are consistent with recent studies where this sensor was instrumental to identify new mutations associated with epilepsy using a high-content screening approach (15).

On another hand, our method can be applied to characterize pharmacological modulators of hNaCT. Both PF-4a and BI01383298, previously described as potent and selective inhibitors, significantly reduced citrate uptake in our system. In contrast, a well-documented enhancer of hNaCT function lithium elicited a reproducible increase in uptake rate. These findings confirm that the system can detect both inhibitory and stimulatory modulation of hNaCT activity, highlighting its potential for drug screening efforts targeting this transporter. Results from single-cell measurement were critical to establish an HTS method that showed stable signal detection of citrate uptake and its pharmacological inhibition, with a Z’-factor above 0.59 across independent experiments. Although our HTS method was performed using a 96-well plate format, sensor-based methodologies have been developed that are readily scalable to 384-well format (16).

Compared to traditional radioactive tracer-based assays, which remain the gold standard for transporter studies, our fluorescence-based method offers several practical advantages. It eliminates the need for radiolabeled compounds, thereby reducing safety and disposal concerns. Moreover, it enables single-cell measurements under standard laboratory conditions, allowing deeper exploration of cell-to-cell variability. Additionally, the method is easily adaptable to HTS format for drug discovery applications.

## AUTHOR INFORMATION

### Authors

**Natali Carcamo-Lemus** – Centro de Estudios Científicos, Av. Arturo Prat 514, postal code 5110466, Valdivia-Chile.

**Angelo Bernier** – Universidad Austral de Chile, Isla teja s/n, postal code 5110566, Valdivia-Chile. **Ivana Oporto-Ortega** – Universidad Austral de Chile, Isla teja s/n, postal code 5110566, Valdivia-Chile.

**Alejandro San Martín** – Facultad de Medicina, Universidad San Sebastián, 5110773, Valdivia, Chile and Centro de Estudios Científicos, Av. Arturo Prat 514, postal code 5110466, Valdivia-Chile.

## Author Contribution

*N.C. and A.B. contributed equally. A.B. and N.C., data analysis. N.C., A.B., and I.O. performed experiments. A.S. reviewed the manuscript and experimental design. P.Y.S. conceived the study, supervised the project, and wrote the manuscript.

## Acknowledgments

We thank the members of the Energy Metabolism Group at CECs for helpful discussions. This work was funded partly by USS-FIN-23-FAPE-03 (PYS), FONDECYT project #11190584 (PYS), and FONDECYT project # 1230145 (LFB).

## Data Availability Statement

Data is available upon request from the corresponding author.

